# Understanding the evolution of interspecies interactions in microbial communities

**DOI:** 10.1101/770156

**Authors:** Florien A. Gorter, Michael Manhart, Martin Ackermann

## Abstract

Microbial communities are complex multi-species assemblages that are characterized by a multitude of interspecies interactions, which can range from mutualism to competition. The overall sign and strength of interspecies interactions have important consequences for emergent community-level properties such as productivity and stability. It is not well understood whether and how interspecies interactions change over evolutionary timescales. Here, we review the empirical evidence that evolution is an important driver of microbial community properties and dynamics on timescales that have traditionally been regarded as purely ecological. Next, we briefly discuss different modelling approaches to study evolution of communities, emphasizing the similarities and differences between evolutionary and ecological perspectives. We then propose a simple conceptual model for the evolution of communities. Specifically, we propose that the evolution of interspecies interactions depends crucially on the spatial structure of the environment. We predict that in well-mixed environments, traits will be selected exclusively for their direct fitness effects, while in spatially structured environments, traits may also be selected for their indirect fitness effects. Selection of indirectly beneficial traits should result in an increase in interaction strength over time, while selection of directly beneficial traits should not have such a systematic effect. We tested our intuitions using a simple quantitative model and found support for our hypotheses. The next step will be to test these hypotheses experimentally and provide input for a more refined version of the model in turn, thus closing the scientific cycle of models and experiments.

## Introduction

Microorganisms play key roles in biogeochemical cycling, industry, and health and disease of humans, animals, and plants[1–5]. For example, the roughly 10_30_ microbial cells on our planet contain ten times more nitrogen than all plants combined and are responsible for half of the global production of O_2_[6]. Almost all of these microorganisms reside in communities, which are assemblages of multiple interacting species. Despite their importance, we currently know little about how microbial communities form and function. However, such knowledge is crucial if we want to fundamentally understand their properties and dynamics, as well as control the processes that they mediate[7,8].

Microbial communities, like all complex systems, are more than the sum of their parts: they are characterized by a multitude of often complex interactions between their constituent members (Figure 1). At any given time, microbes may compete for shared resources such as metabolites and space, inhibit each other via the secretion of antibiotics and other toxic compounds, and even kill each other upon direct cell-cell contact[9,10]. Yet, not all is bleak in the microbial world: some organisms may — accidentally or actively— excrete enzymes or molecules that others can use, and even commit suicide for others [11–13]. The overall sign and strength of an interaction between two organisms is the net result of all such processes and can be characterised as anything from competition to parasitism and mutualism[14,15].

**Figure 1.**
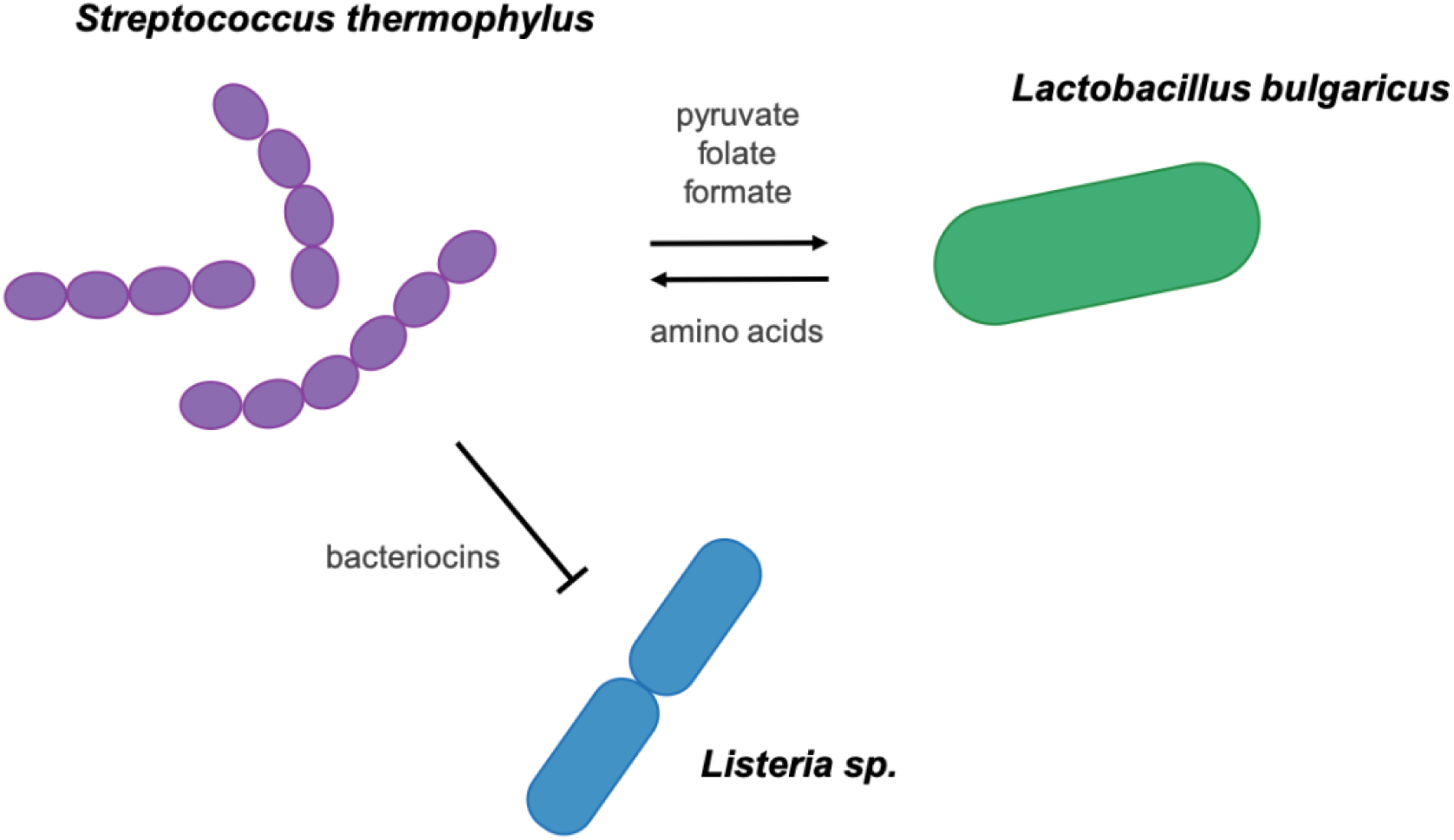
Microbial communities are characterized by a multitude of often complex interactions between their members. For example, three bacterial species commonly found in yoghurt cultures engage in both positive and negative interactions, which are mediated by metabolic compounds and toxins[19,20].

Interspecific interactions have a crucial role to play in microbial communities. That is, what makes a community a community— rather than a random set of species— is precisely these interactions, because they give rise to properties at the level of the community that we cannot understand by considering each species in isolation. For example, it may not be possible to predict from growing each species by itself what the joint reproductive output of a collective will be, or how robust such a collective will be to external biotic and abiotic perturbations. Microbial communities can also perform chemical transformations that would be impossible for one individual species to achieve[16], and some communities even display complex behaviours such as collective motion and electrochemical signalling, which have traditionally been associated with higher organisms[17,18].

Over the past decades, an impressive amount of thought and effort has been dedicated towards obtaining an ever-more detailed and realistic picture of a wide range of different microbial systems. The advance of omics technologies has led to an incredible leap forward in terms of available data that can be integrated to extract and analyse patterns that inform us about the lives of microbes in their natural surroundings. However, if we want to arrive at a more fundamental understanding of the current and future properties of microbial communities, we will have to uncover general principles of how such communities will typically change over time.

Since interactions are often mediated by whole suites of different chemicals, interactions may significantly change in strength and even sign over ecological timescales. For example, bacteria may alter the pH of their environment during growth, which may then change the sign of their interactions with other species from positive to negative, or *vice versa*[21]. Extrapolating such findings to more general principles of community dynamics is challenging. Nonetheless, an increasing number of studies finds that ecological community dynamics are remarkably repeatable across experimental and biological replicates, and may be understood from a combination of metabolic properties of the environment and species functional traits[22–24]. Less is known about how communities change over evolutionary timescales, even if such changes have potentially much more far-reaching consequences because they are essentially irreversible (i.e. they are the result of mutations, so the community will not change back to its original state unless these mutations are reverted, or species acquires additional ones). Given that interactions are the defining feature of a community, we propose that what is most important in this respect, is how interspecific interactions change.

Here, we review the empirical evidence that evolution is an important driver of microbial community properties and dynamics on timescales that have traditionally been regarded as purely ecological. Next, we briefly discuss different modelling approaches to this problem, emphasizing the similarities and differences between evolutionary and ecological perspectives. We then propose a simple conceptual model for the evolution of communities, which we explore using simulations. Finally, we discuss experimental approaches that may help to test our framework and thus improve our understanding of this fascinating process.

## Evolution of microbial communities: empirical approaches

The question of how microbial communities evolve has been addressed empirically by authors from several different fields. One approach that has substantially increased in popularity over recent years is genomic analysis of one or more— often pathogenic— species that evolve within a host environment, such as the gut or the cystic fibrosis lung[25]. Some of these studies employ temporal metagenomics to capture the entire diversity present within a given population, whereas others use culture-based population genomics to obtain more reliable estimates of the phylogenetic relationship between genetic variants[26,27]. This group of studies has revealed that evolution in the microbiome can be fast and highly repeatable[28], with 109 to 1012 new single nucleotide polymorphisms arising on a daily basis[27], *de novo* mutations competing for extended periods of time[29], and gene gains and losses sweeping to high frequencies within a period of just a few months[30].

Metagenomic approaches have also been used to study evolution in microbial communities that are not host-associated. A nine-year study of biofilm communities from acid mine drainage allowed the authors to estimate the *in situ* nucleotide substitution rate for a focal species and revealed several divergence and hybridisation events[31]. A similar study performed on a freshwater lake community monitored multiple species simultaneously and found evidence for extensive within-population genetic heterogeneity, as well as selective sweeps in some species but not others[32]. Horizontal gene transfer also appears to be an important driver of adaptation in natural communities, at least when they are exposed to selective pressures such as antibiotics or heavy metals[33,34]. Finally, metagenomic approaches have been used in conjunction with phylogenetic tools to infer evolutionary rates of microbial communities as a whole over longer timescales[35]. These studies underline the importance of the environment for the rate of adaptation, with communities evolving most slowly in energy-limited environments that impede growth[36] and evolving most rapidly in extreme environments that impose strong selection[37].

Another field that is concerned with the evolution of microbial communities, albeit from a more applied angle, is artificial community selection. This approach entails the repeated selection of microbial communities from a larger pool based on their performance with respect to some *a priori* defined community-level function, such as productivity or enzyme production[38–40]. Given that this approach does not usually assess genetic changes in each of the constituent species, it is conceivable that the observed changes are mainly due to ecological sorting rather than evolutionary adaptation[41,42]. However, the duration of these experiments is sufficient for evolution to occur, and it has in fact been argued that continued species coexistence and within-species evolution are crucial for maximally-effective community selection[43]. One important advantage of this approach over methods that are centred on focal species is that it explicitly tracks the dynamics of the entire community. As such, the success or failure of this approach may provide valuable insights into whether we can regard communities as a unit of selection[44–46] and to what extent we can predict changes in community-level properties over time.

A more bottom-up approach to studying the evolution of microbial communities is laboratory evolution. This method involves culturing replicate populations of organisms under controlled conditions for long periods of time, while probing the resulting phenotypic and genotypic changes[47,48]. While this method has been used mainly to look at the evolution of single species, an increasing number of studies employs simple two-species communities to ask how species evolve in the presence of a coevolutionary partner[49]. Mostly, these experiments focus on the evolution of species with a predefined interspecific relationship— such as host-parasite[50], predator-prey[51], or mutualism[52,53]— and assess how interaction traits like parasite infectivity and host defence change over evolutionary time. At least two studies explicitly report a change in the nature of the interactions between two species: in one case, a commensal interaction quickly evolved into exploitation[54], whereas in the other case, ammensalism evolved into antagonism[55].

Given the increasing popularity of laboratory evolution as a tool and the fact that natural communities often consist of a large number of interacting species, surprisingly few studies have extended the above approaches to communities of more than two species (but note Refs. [56,57]). One important exception is a study in which five species from a beech tree hole were cocultured in the lab for ∼70 generations[58]. Characterisation of the evolved communities revealed that species had diverged in resource use and evolved to feed on by-products excreted by the other species. As a result, interspecific interactions became more positive (in the sense that growing species in pairs lead to a higher yield than would be predicted based on growing them individually), and community productivity increased. The same qualitative results were also recovered when between 1 and 12 species from the same model community were evolved under three different conditions[59], again demonstrating the importance of studying the evolution *of* microbial communities, rather than evolution *in* microbial communities.

Taken together, the above approaches provide us with a wealth of data on the evolution of microbial communities, which gives us a first rough understanding of the nature of this process. The emerging picture is that evolution is likely to be an important driver of community dynamics and properties on what have traditionally been regarded as ecological timescales[60–62]. One major scientific challenge—and opportunity—is now to develop a framework that allows us to *predict* evolutionary changes in microbial communities, based on theoretical and mechanistic principles. Our goal here is to contribute to the development of such a framework.

## Evolution of microbial communities: modelling approaches

To test our intuitions about general principles of community evolution, one powerful approach is to use mathematical and computational models. Such models allow us to assess how features of a system interact, ideally generating testable hypotheses that can be addressed using carefully designed experiments[63,64]. In particular, they allow us to test a wider range of possibilities, such as environmental conditions or combinations of species, than would be experimentally feasible.

There are many models describing aspects of the evolution of microbial communities, but few of these models consider the process in its entirety. This is maybe not surprising, given that community evolution is a complex problem that requires an understanding of microbiology as well as a combination of modelling approaches from evolution and ecology, two disciplines that have traditionally been poorly integrated despite their conceptual similarities. However, comparing approaches from both fields and combining the relevant elements from each of them may be a promising tactic towards tackling this problem. To this end, we here briefly review the main properties of some common evolutionary and ecological models and evaluate their usefulness for understanding the evolution of microbial communities.

### Evolutionary models

One important class of evolutionary models is population genetic models. These models are focused on tracking changes in genotype or allele frequencies over long timescales (Figure 2)[65]. Different genotypes arise via mutations or migration and map to some measure of reproductive success, or *fitness*[66], perhaps via an intermediate phenotype. The genotype with the highest fitness increases in frequency until it eventually replaces all other types or is outcompeted by new mutations with an even higher fitness[67]. Many variations on this theme exist with added levels of complexity, such as recombination[68], spatial structure[69], and more complex demography[70].

**Figure 2.**
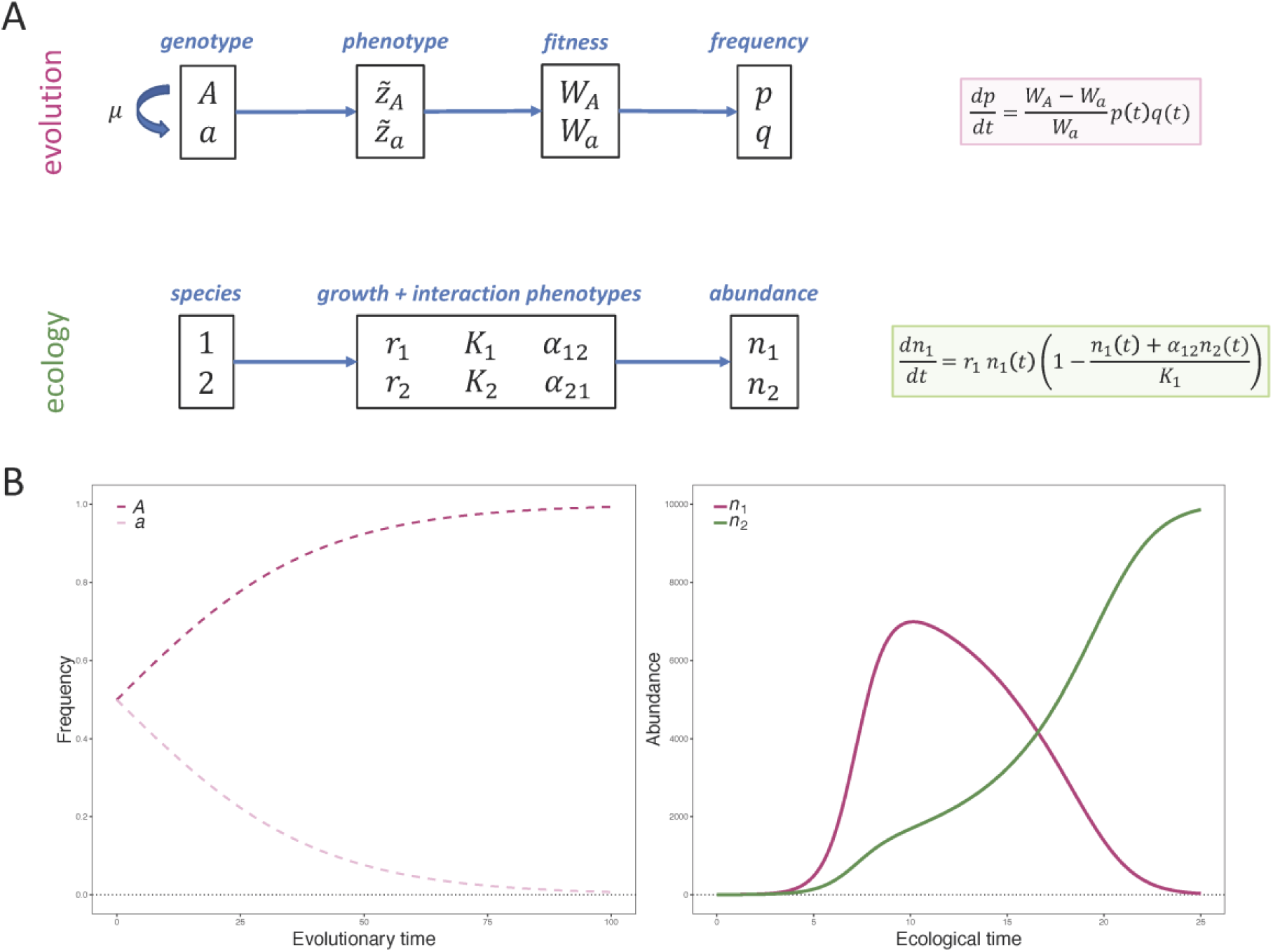
Conceptual set-up of classical models in evolution and ecology. A) Population genetics models consider different genotypes that can interconvert via mutation (*μ*). Each genotype maps to some fitness (*W*) value, perhaps through an intermediate phenotype. A mathematical model (pink box) then describes the change in frequencies of the genotypes (***p*** and ***q***) over long timescales, where the average speed of this process depends on the relative fitness of each type. In contrast, ecological models consider multiple species, each of which maps to a set of growth and interaction phenotypes (*r, K, α*). A mathematical model (green box) tracks the changes in abundances of different species (*n*_1_and *n*_1_) over shorter timescales. B) Left: dynamics of genotype frequencies from the evolutionary model (*p*_t0_= *q*_t0_= 0.5, *W*_*A*_= 1.05, *W*_*a*_= 1.00). Right: dynamics of species abundances from the ecological model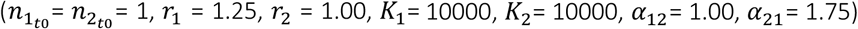.

In contrast, quantitative genetic models typically focus on traits encoded by a large number of loci, which are assumed to evolve independently due to substantial recombination, as is common in animals and plants [71]. Therefore, these models ignore the underlying genetics of the phenotype and instead track the change in the mean phenotype 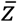 of the population over time. Populations are composed of many different types that each have a different deviation 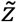 from the mean, as well as a different fitness. Selection then acts on this variation to increase population fitness. The overall relationship between phenotype and fitness is crucial: larger total variation in phenotype and larger selection on that phenotype lead to faster evolutionary change [72,73].

Because evolutionary models typically consider dynamics over long timescales, the model parameters such as fitness or population sizes are usually some average values that integrate over many underlying phenomena. For example, the fitness may be an average fitness over many possible environments that fluctuate among each other or over many stages of life history. As a result, it can be difficult to link these models to explicit biological mechanisms. In particular, explicit interactions between organisms and their biotic and abiotic environment are almost always neglected in these models. The reason is that the dynamics of these interactions are assumed to be much faster than evolutionary dynamics and hence can be averaged out. However, as aforementioned, it is increasingly clear that ecological and evolutionary dynamics often occur on similar timescales in microbes. Therefore, an explicit treatment of ecological processes should be necessary to more accurately describe evolution.

### Ecological models

Arguably the most classic ecological model is the Lotka-Volterra model (Figure 2)[74]. This model tracks over time the abundance of two or more different species, which are assumed to be immutable. When the species are growing far from their *carrying capacity K*, or maximum population size at which the population has a positive growth rate, the abundances increase exponentially and independently between species. However, as the population sizes increase, intraspecific and interspecific interactions reduce growth, such that the species with the fastest maximum growth rate may not dominate in the long run. An advantage of ecological models is that they make explicit possible trade-offs between life-history traits: for example, when organisms are growing in quickly changing, resource-rich environments far from carrying capacity (e.g., frequent migration into new resource patches), it pays off to grow faster, but when growing in constant, resource-poor environments close to carrying capacity (e.g. biofilms), it pays off to be more efficient or have a more inhibiting effect on your competitor.

The Lotka-Volterra model is purely phenomenological, i.e. it describes the dynamics of the process without making any direct reference to the underlying mechanisms[74]. By contrast, consumer-resource models explicitly track the resources that are responsible for the observed dynamics[75]. Such added level of detail can help to understand why we observe certain dynamics and make predictions beyond the current observations. It is even possible for phenomenological and mechanistic models to generate opposite predictions about the dynamics of the system for relatively simple cases[76]. However, for each additional quantity that is tracked over time, additional parameters are needed, and in the case of microbial communities, where many chemically-mediated interactions occur simultaneously, keeping track of and explicitly modelling each of these interactions is daunting.

One approach that can potentially overcome this problem is community-level metabolic modelling known as flux balance analysis (FBA), where the metabolic conversions performed by each species in the community are predicted from the species’ genomes, under the assumption that each cell maximises its growth rate (i.e., the rate of biomass production)[77]. While simpler versions of these models assume that each of the individual species are in steady state, more complex versions allow for variation in metabolic flux depending on the environment, and thus the tracking of the abundance of each individual species over time [8].

A second important extension of these models is the incorporation of spatial structure[78]. Given that most microbes live in spatially-structured environments[79] and chemicals can only travel limited distances[80], it is clear that microbes do not usually interact equally with everyone else in the population, but instead interact most strongly with their immediate neighbours[11]. As such, growth rate as well as interaction parameters such as *K* and *α* will vary over space and time. This may lead to locally different outcomes of competition, which can have important consequences for the reproductive success of each type within the population as a whole. Such local variations in population dynamics and their global consequences are commonly explored using individual-based simulations[81,82], where each cell grows, dies, and moves depending on its local context. The advantage of such approaches is that they can explore the emergent properties of complex systems where many factors interact, especially for cases where analytical solutions to the problem cannot be obtained.

### Integrating evolution and ecology

Considering the classes of models reviewed above, it is clear that for a model to adequately capture the evolution of microbial communities, it will have to incorporate elements from both evolution and ecology. While some processes, such as mutation or kin selection[83], are particular to within-species dynamics, many other elements from both disciplines can in principle be used interchangeably, as the distinction between different genotypes of the same species and different species is often arbitrary. Specifically, a model suiting our purpose would have to allow for mutation and selection, as well as intra- and interspecific interactions. Additionally, because interactions are often local and mediated by chemicals, spatial and mechanistic models may be more accurate in many situations compared to mass-action and phenomenological models. Finally, for a model to capture the evolutionary dynamics of the community as a whole, it will have to allow not only for the evolution of each of the individual species, but also for the evolution of the interactions between them.

Some models already exist at this intersection of evolution and ecology. One emerging discipline is eco-evolutionary dynamics, which focuses on the interplay between the composition of species and the abundance of these species[60,62,84,85]. However, the most notable category of interest for the current problem is evolutionary game theory[86–89], where the fitness of a phenotype depends on the frequency of the other phenotypes in the population. The mathematical description of these models is very similar to that of the Lotka-Volterra equation[90], even if frequency- and density-dependence do not necessarily have the same consequences (consider, for example, the time points in Figure 2B where the two types have equal frequencies but different abundances). These models are often used to investigate how costly cooperative traits within species (the production of “public goods”), such as the secretion of extracellular enzymes that catalyse the transformation of resources[91] or toxins that kill a competing strain or species[92,93], can evolve or be maintained in the presence of defecting or cheating individuals that do not pay the cost but gain the benefit. Sometimes they also incorporate mutation[94] to explore how the mean trait level within a species evolves over time. This approach has also been applied to the evolution of interspecific interactions[95,96], although the underlying traits are often different in that case[97].

### Evolution of interspecific interactions

Most modelling approaches that assess how interspecific interactions change over evolutionary timescales focus on one specific type of interaction, such as interference competition[98] (where two species interact in a way that is detrimental for both species, an interaction that can be represented as “-/-”), host-parasite[99,100] or predator-prey interactions[101] (+/-), or mutualism[102,103] (+/+). Generally, these models look at how a single trait such as parasite infectivity or host defence changes over time by considering different types (which may or may not be generated by mutation) within each species, how their frequencies change over time, and what consequences this has for the population as a whole.

Understanding how such traits evolve is important when studying the evolution of microbial communities. However, not all microbial interactions can be classified with such simple schemes, and the nature and strength of ecological interactions often changes with time[21,104,105]. Moreover, even when the nature of an interaction is stable, this does not necessarily mean that the phenotypic traits on which the interaction is based will evolve, because species may adapt to their local biotic and abiotic environment in myriad other ways, particularly when the community consists of more than two species. So the question is not whether such interspecific interactions can evolve, but what the likelihood relative to other traits is that they will evolve.

Evolutionary changes in interspecies interactions are expected to strongly depend on the underlying costs and benefits. Exerting positive as well as negative effects on other species is often costly. For example, the excretion of compounds that increase another species’ growth— e.g., enzymes, building blocks, metabolites, or chelators— is expected to exert a metabolic burden on the producer[12,106,107]. The same is true for compounds that have inhibitory effects on other species, for example, antibiotics and other antimicrobial compounds or detrimental effectors that are delivered in a contact-dependent way[9].

Given these metabolic costs, why do so many microbes invest resources in affecting other species in positive or negative ways? One explanation is that these effects on others can lead to indirect benefits. Increasing the growth and activity of another species can be beneficial if this other species is a mutualistic partner. Similarly, suppressing growth and survival of a species can be beneficial if this other species is a competitor. If we want to understand the evolution of interspecies interactions, we thus need to understand how such indirect benefits arise, which individuals have access to these indirect benefits, and how the indirect benefits compare to the costs of investing into an interaction.

## Conceptual and computational models for the evolution of microbial communities

Our next goal is thus to consider the emergence of indirect benefits that arise from an evolutionary change of an interspecies interaction. One central idea is that mutants that invest more into an interspecies interaction – be it mutualistic or antagonistic – can only increase in frequency if they have preferential access to the indirect benefits they create[95–108,114]. To illustrate this, consider the fate of a mutant that produces a costly antibiotic that kills a competing species. When such a mutant arises in a well-mixed environment, such as a shaking flask, all its conspecifics will now be able to grow faster because their competitor is inhibited, while they do not pay the cost of antibiotic production. As a consequence, they will grow more rapidly than the mutant. The mutant will thus not be able to increase in frequency within its own population. By contrast, when the same mutant arises in a spatially-structured environment, such as a biofilm, the mutant and its descendants will profit more than their average conspecific from the inhibition of their neighbouring competitors. This is because the inhibitory compounds, as well as the extra nutrients that are now available, are locally more concentrated. In spatially-structured environments, such a mutant is expected to increase in frequency within its population, given that the associated cost is not too high.

Based on the above argument, we hypothesize that evolution in well-mixed environments will proceed exclusively via the selection of directly-beneficial traits. By contrast, evolution in spatially-structured environments may proceed via the selection of both directly- and indirectly-beneficial traits (Figure 3A); which of these two modes predominates will depend on the availability and effect size of mutations on each type of trait. Importantly, mutations that provide a direct fitness benefit may also affect other organisms in both positive and negative ways as a pleiotropic effect. For example, when an organism becomes better at using a resource and collaterally produces more metabolic by-products, others may also profit from this. However, such a mutation will not be selected because of the effect that it has on others, but because of the effect that it has on the actor itself.

**Figure 3.**
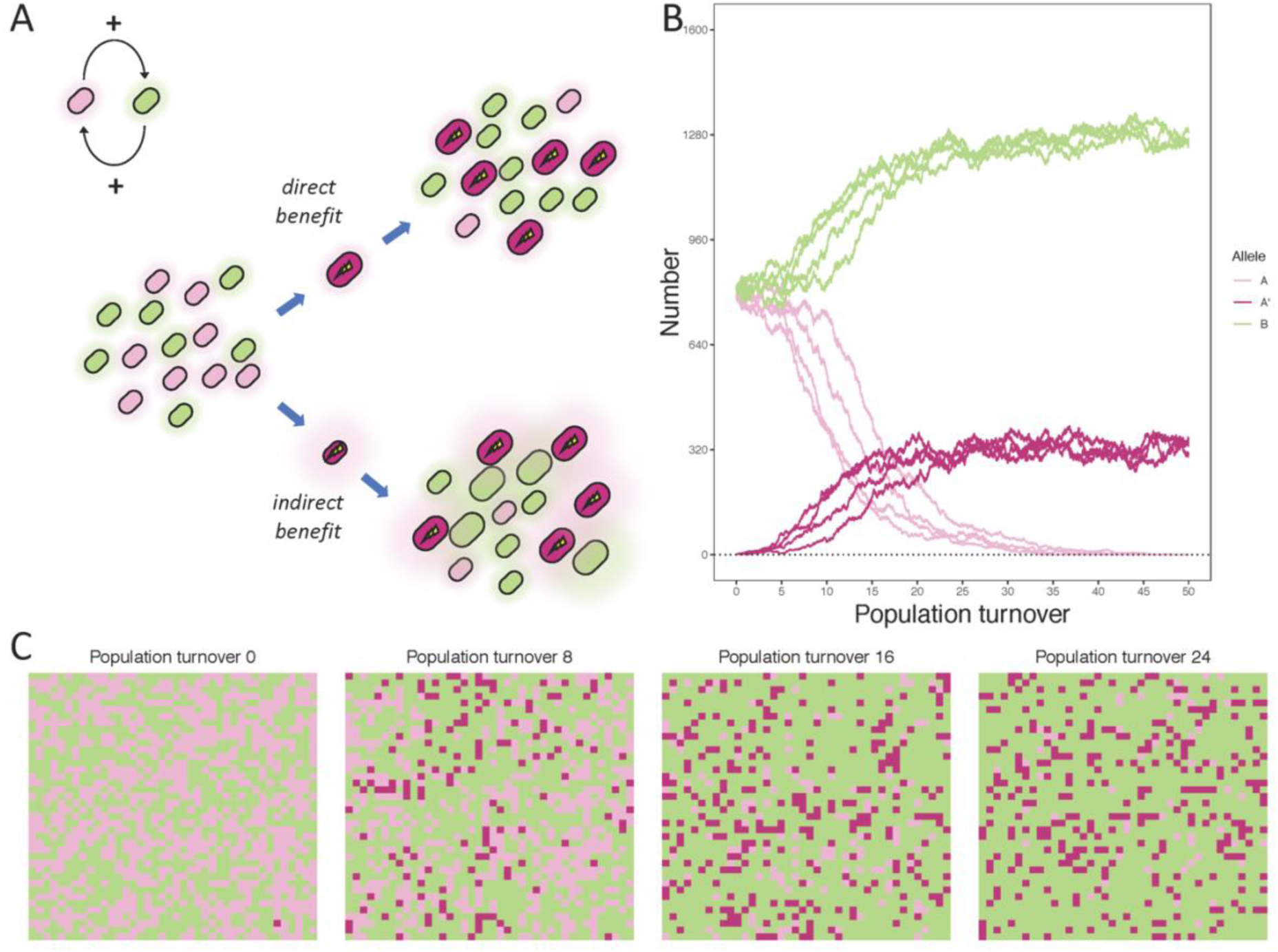
A conceptual and computational model for the evolution of microbial communities. A) Schematic depicting the hypothesis put forward in this paper. We here illustrate the case of two species (pink and green) having a positive initial effect on each other, but the same principle applies to other types of initial interactions. Pink individuals can evolve by the acquisition of two types of mutations. First, they may acquire mutations that provide a direct fitness benefit, such as becoming better at using a certain resource, or becoming more tolerant to a toxic compound. Individuals carrying such mutations should be able to increase in frequency in both spatially structured and well-mixed environments. Second, they may acquire mutations that provide an indirect fitness benefit, such as producing a costly compound that increases the growth of the green species. Because the green species has a positive effect on the pink species, having more green cells around, as well as each of these cells having a higher growth rate, also provides a fitness benefit to the pink species (e.g. because the green cells excrete a metabolic by-product, that the pink species can use, at a rate proportional to their growth rate). Individuals carrying such mutations should only be able to increase in frequency in spatially structured environments. Cell size reflects growth rate. B) Population dynamics in 10 replicate runs of a simple cellular automaton model, simulating a spatially structured environment. Alleles A and B grow better when there are more cells of the other type in its immediate neighbourhood. A costly, more cooperative A’ mutant invaded in 4/10 simulations and reached an intermediate equilibrium frequency. By contrast, this mutant invaded in 0/10 replicates of a model simulating a well-mixed environment. C) Spatial dynamics in one replicate run of the model describes in B.

To test the idea that spatial structure will determine whether mutations with indirect benefits are selected for, we developed a simple cellular automaton model[92] that incorporates elements from both evolution and ecology: it allows for intra- and interspecific interactions as well as the possibility of these interactions evolving by mutations. We assume that cells of two different species, A and B, grow together on a 40×40 grid with periodic boundary conditions. The fitness of each cell depends on the fraction of cells around it of each type of species, as well as the fitness of each of these neighbouring cells. So, an interaction impacts both the reproductive success of the other species as well as its reciprocal interaction. In each time step, one cell in the grid reproduces with a probability proportional to its fitness. We then randomly choose another cell to die, with the descendant of the reproducing cell pushing all cells in its path towards the now-empty spot vacated by the dead cell. A complete population turnover will take on average 40×40 = 1600 such time steps. For details of the model, see the Methods section.

Here we focus on the scenario where both types already have a slightly positive effect on each other (i.e., an example of passive mutualism[108]). We assess whether a single mutant A’, which has a more positive effect on B but grows more slowly than the other A cells, can increase in frequency within the population. Note that, even if we do not explicitly test this here, we predict that the same logic will also apply to the case of where both species have a slightly negative effect on each other and a costly mutant with a stronger negative effect emerges. We run the simulation for 50 whole population turnovers and determine the probability that the mutant establishes, i.e. rises above some minimum abundance so that it is unlikely to be lost by stochastic effects. In parallel, we run control simulations for a well-mixed environment, where fitness of a cell depends not just on its neighbours, but on all other cells in the population.

For the parameter settings that we chose, we find that a costly mutant with a strong positive effect on a cooperating species invades in 41 out of 100 cases (Figure 3B, Methods). When it does, the original A genotype is driven to extinction, and the mutant reaches an equilibrium concentration of ∼20% (that is, A’ equilibrates at a lower frequency than the other type, B, presumably because it invests into an interaction that is beneficial for B). By contrast, in the well-mixed environment, the mutant invades in only 2 out of 100 cases, and when it does, it reaches only very low, fluctuating concentrations, suggesting that such invasion is the result of genetic drift. While the precise dynamics depend on the details of the model as well as the parameter settings[115], our goal here is to provide a proof of principle showing that mutations that provide an indirect fitness benefit can be selected for in a multispecies community.

An interesting future extension of such models is to allow for mutants providing direct and indirect fitness benefits to arise simultaneously, and to investigate how the success of such mutants depends on the composition of each species as well as their abundances. For example, based on the models discussed above, we might predict that mutations with a direct fitness benefit will have a relative advantage at lower densities, because they increase growth rate (*r*), whereas mutations with an indirect fitness benefit will have a relative advantage at higher densities, because they impact interactions (*α*). Another interesting extension is to investigate the impact of genetic architecture and the correlation structure of mutational effects on self *vs.* non-self. That is, are mutations with a direct *versus* indirect fitness benefit more likely to occur, and/or have a different distribution of fitness effects? Does a mutation with a more positive effect on the actor generally have a more positive effect on others around it as well? Finally, it might be insightful to explicitly model the different classes of metabolites that are expected to underlie the interactions and to assess whether such a mechanistic approach makes predictions that are qualitatively different from those of the phenomenological approach that we used here.

The generation of a conceptual, qualitative model— in which reality is abstracted away to its minimal essence (Figure 3A)— and a simple, quantitative model— to see whether our intuitions bear out at least in principle (Figure 3B)— are useful first steps towards understanding the evolution of microbial communities. However, science is ideally an iterative process of models and experiments, where each informs the other. One way to test whether our hypotheses can stand the test of reality is to perform evolution experiments with multiple microbial species in both spatially-structured and well-mixed environments. Careful phenotypic, genotypic and metabolic characterisation can then be used to disentangle mutations with direct and indirect fitness benefits.

## Scaling up

Whether traits typically confer a direct or indirect fitness benefit to the actor can have important consequences. When a mutation in a focal species gets selected for its effect on a second species, be it positive or negative, this second species will now be under stronger selection to acquire a mutation with a similar effect on its partner as well. This is because the indirect benefit that the second species can obtain from an increased investment into the interaction has increased as well. As such we would predict that when evolution proceeds predominantly via the selection of mutations with an indirect fitness benefit, interaction strength will increase over time, potentially to the point of obligate mutualism or bi-stable exclusion. When species initially have an opposite effect on each other (+/-), the long-term direction of this coevolutionary cycle is less clear. However, one possibility is that the outcome depends on which species mutates first. For example, when a first species evolves to excrete more of a compound that inhibits a second species, which has a positive effect on the first species, this second species now will be under stronger selection to decrease its help to the first species, or even to start harming it. Clearly, this prediction will not hold for interactions of which the sign cannot change by definition, such as host-parasite interactions (but note that even such interactions may not be constant[116,117]). Nonetheless, microbial communities are dominated by chemically-mediated interactions, suggesting that the selection of indirectly-beneficial mutations may occur also in the case of asymmetric interactions.

The overall sign and strength of interspecific interactions have important consequences for properties at the level of the community. More positive interactions are associated with increased productivity, because fewer resources are wasted on competitive traits, and species may engage in division of labour[118,119]. At the same time, an overrepresentation of positive interactions will make a community less stable, because it decreases the number of negative feedback loops, and consequently, when one species is perturbed, it may drag the others down with it to extinction[120]. By contrast, more negative interactions should have a stabilising effect on the community, because any interaction loop with an odd number of negative connections will result in a negative feedback loop[121]. Negative interactions will also decrease productivity, and whenever interspecific competition becomes stronger than intraspecific competition, this may lead to bi-stable exclusion[122]. More generally, the topology and interaction strengths of any type of network, including the interaction network of a microbial community, are predicted to have far-reaching consequences for the stability and resilience of the system[123–125].

One intriguing question that has received an increasing amount of interest over the past few years is whether microbial communities may be regarded as coherent units, on which selection can act directly [126,127]. If this were the case, we might be able to select for certain desired community-level properties, such as efficiency in producing a specific compound[43]. While this idea has a strong intuitive appeal, experimental results have been mixed[128]. This is most likely due to the fact that for selection to act on something, a minimum amount of heritability is needed: if a microbial community breaks up into its component parts, or changes in composition due to ecological and evolutionary forces, sufficiently often relative to it giving birth to a similar community, natural and artificial selection will not be able to act upon the properties of the community.

One way in which such heritability may be increased is if species become so strongly dependent on each other as a result of coevolution that they can no longer live alone[129]. Alternatively, it may result from organisms having coevolved in a community context for sufficiently long that each of them has become locally adapted to its biotic environment, such that foreign species will be unable to replace their resident counterparts[44,130].

Interestingly, our hypotheses predict that both of these types of coevolution should occur significantly more often in spatially-structured environments (because it is there that mutations are selected for their indirect fitness effects), which might be an interesting idea to test experimentally. Increased metabolic dependency may also change the spatial association between organisms itself, suggesting that this process may even be subject to a self-reinforcing feedback loop[108].

## Conclusions

The evolution of microbial communities is a fascinating and complex process that has potentially far-reaching consequences. To understand it, we need an integrated approach that encompasses elements from microbiology, ecology, and evolution. Here, we reviewed the evidence for the importance of the evolution of microbial communities in host-associated as well as free-living communities, and integrated previous empirical and theoretical findings to arrive at a conceptual model of this process. We tested our intuitions using a simple quantitative model and found support for our hypotheses. The next step will be to test these hypotheses experimentally and provide input for a more refined version of the model in turn, thus closing the scientific cycle of models and experiments.

## Methods

In our cellular automaton model, we simulated the cell division and death dynamics of two species, A and B, on a 40×40 grid with periodic boundary conditions, mimicking a spatially-structured environment. The A species also has a mutant variant A’. The fitness of each cell *i*, which determines the probability it will divide at the next time step, is the sum of a basal fitness *r* and a term proportional to the total fitness of all cells of opposite species in its immediate eight-cell neighbourhood, weighed by an interaction parameter (*v*):

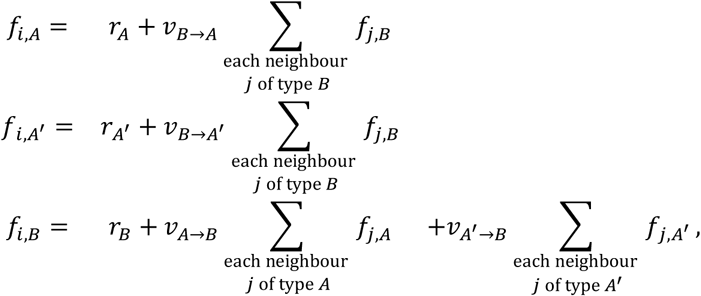

where the basal fitness paramete rs are *r*_*A*_= *r*_*B*_ = 1 and *r*_*A*′_ = 0.9, while the interaction parameters are *v*_*B*→*A*_ = *v*_*B*→*A*′_ = *v*_*A*→*B*_ = 1 and *v*_*A*′→*B*_ = 5. That is, the wild-type A and B cells have identical basal fitness and help each other equally, while the mutant A’ provides a stronger benefit to B but at a cost to its basal fitness.

We start with a grid that is fully occupied by cells randomly chosen to be either A or B, each with initial fitness randomly drawn from a uniform distribution. In each time step, we start by updating the fitness of all cells according to the above equations, constraining growth rates between 0 and 1. Then we randomly choose one cell in the grid to reproduce with probability that is proportional to its updated fitness, placing the new cell in one of the eight neighbouring positions. Next, we randomly choose another cell on the grid to die, and the new cell pushes all cells in the direction in which it is placed relative to its mother cell towards the now-empty spot, taking a path with only a single turn. We define one population turnover as 40 ×40 =1600 of such timesteps.

We started by running this simulation for 10 population turnovers to let the population reach an approximately steady-state configuration. Then we introduced a single A’ mutant and ran the simulation until either the mutant went extinct or reached an “establishment” threshold, which we defined as 10 cells; we chose this threshold to be twice the maximum number of mutant cells that we observed for a mutant destined for extinction (based on 20 preliminary trials). We repeated this procedure 100 times, all starting from the same initial population. Altogether the mutant established in 41 out of 100 of these simulations.

As a control, we performed the same simulations but let the fitness of each cell depend on interactions from all other cells in the population (rather than just cells in the immediate neighbourhood), thus mimicking a well-mixed environment:

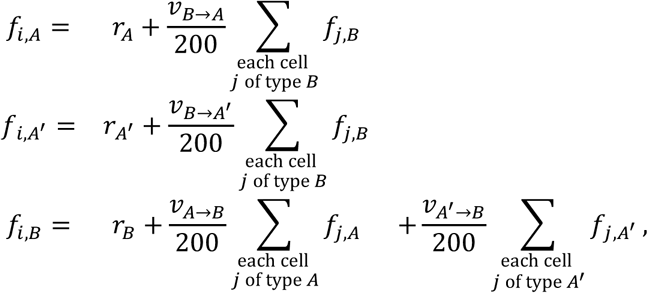

where we rescale the interactions by 200 to keep parameters comparable between the two models, since in the well-mixed case each cell has approximately 200 times more neighbours than in the spatially-structured case.

We chose 16 cells as the establishment threshold in this case, since 8 cells was the maximum number reached by mutants destined for extinction (based on 20 preliminary trials). Based on this criterion, we observed establishment in 8 out of 100 simulations. Because we suspected that these putative establishments might actually be due to genetic drift rather than positive selection, we continued these simulations for another 50 population turnovers. Six of these cases indeed led to extinction during that time, while in the remaining 2 cases, the mutant was still present at low, fluctuating frequencies (<4%) by the end and did not substantially affect the frequency of the wild-type A.

We plotted the first 10 replicates of each simulation for Figure 3. We ran all simulations in R.

## Acknowledgements

FAG was supported by an ETH Fellowship and a European Molecular Biology Organization (EMBO) Long-Term Postdoctoral Fellowship (ALTF 335-2018); MM was supported by an Ambizione Fellowship from the Swiss National Science Foundation (PZ00P3_180147); FAG and MA were supported by the Simons Foundation: The Simons Collaboration on Principles of Microbial Ecosystems (PriME #542389).

